# Validation that human microbiome phages use alternative genetic coding with TAG stop read as Q

**DOI:** 10.1101/2022.01.06.475225

**Authors:** Samantha L. Peters, Adair L. Borges, Richard J. Giannone, Michael J. Morowitz, Jillian F. Banfield, Robert L. Hettich

## Abstract

Metagenomic findings suggesting that bacteriophages (phages) can use genetic codes different from those of their host bacteria reveal a new dimension of phage-host interaction dynamics. Whereas reassignment of stop codons to code for amino acids has been predicted, there has been no proteomic validation of alternative coding in phages. In fact, one code where the stop codon TAG is reassigned to glutamine (code 15) has never been experimentally validated in any biological system. Here, we characterized stop codon reassignment in two crAss-like phages found in the human gut microbiome using LC-MS/MS-based metaproteomics. The proteome data from several phage structural proteins clearly demonstrates reassignment of the TAG stop codon to glutamine, establishing for the first time the expression of genetic code 15.

**One-Sentence Summary:** Mass spectrometry confirms protein expression of predicted alternate genetic coding in phage genomes from human microbiomes.

## Main Text

Bacteriophages modulate the composition of microbial communities through the selective predation of bacteria, alteration of host metabolism, and redistribution of cellular lysis products in the environment during the infection process(*1*). Despite their recognized importance as components of ecosystem dynamics, phages remain one of the most poorly understood members of microbiomes(*2, 3*) due to limitations of the methodologies used to study them. Fundamental questions remain regarding how phages interact with, and redirect, the translation systems of their host bacteria.

There is evidence from metagenomic studies that some phages appear to use the bacterial ribosome to translate their proteins using both the standard and an alternative code. In these phages, proteins that require a stop codon to be read as an amino acid are thought to be only translated late in the infection cycle after code switch machinery has been deployed(*4*). Some phages are predicted to reassign the normal stop codon TAG to be translated as glutamine (Q), and others reassign the TGA stop codon to tryptophan (W). This phenomenon appears to be common in human and animal microbiomes(*1, 4–8*) and is particularly prevalent in phages that infect Firmicutes and Bacteroidetes(*4*).

If not recognized, stop codon reassignment can limit our ability to identify phages in metagenome sequences and restrict our understanding of phage gene inventories. Specifically, incorrect code usage leads to low predicted coding densities, truncated gene products, and genes predicted in incorrect reading frames. Alternative code choices can be tested to determine if they restore full-length open reading frames, and the amino acid to which a stop codon is reassigned can be predicted based on amino acid alignments with homologous proteins from related phages. Direct proteomic confirmation of alternative coding predictions has been primarily restricted to bacteria(*9, 10*), and alternative coding predictions in phages have never been validated experimentally. Bioinformatic studies have inferred the use of genetic code 15 in some bacteriophages(*4, 5, 8*) and the protist *Iotanema spirale*(*11*). However, there are no reports validating the expression of this alternate code, nor is it listed in the summary of genetic codes recognized by NCBI (https://www.ncbi.nlm.nih.gov/Taxonomy/Utils/wprintgc.cgi).

Our previous metagenomic studies identified two unrelated human microbiome samples (adult and infant) that each contain an abundant crAss-like phage that appear to use genetic code 15. The adult sample contained a 191 kilobase crAss-like phage genome with the potential to circularize, and the infant sample had a 94 kilobase crAss-like phage genome, which was curated to completion (**Fig. S1**). These samples were prioritized for metaproteomic measurements to address two key questions: 1) can proteins of phages be detected in the presence of highly abundant bacterial, human, and dietary proteins, and 2) can phage proteins be detected that confirm the expression of alternative genetic code 15?

To answer these questions, paired metagenomic and metaproteomic measurements were conducted on fecal samples from one infant and one adult, where metagenomic data indicated an abundance of alternatively coded phages. To ensure accurate peptide identifications from the metaproteomes, assembled metagenomic data from the same samples were used to generate databases of phage proteins that were predicted using either the standard genetic code 11 (TAG→stop) or alternative genetic code 15 (TAG→Q). Phages contribute a relatively small proportion of proteinaceous biomass in fecal samples, making detecting their proteins by shotgun proteomics particularly challenging. In fact, initial measurements of the fecal samples detected no phage proteins. Thus, a combination of centrifugation and filtration-based enrichment techniques was employed to enrich phage particles and their proteins irrespective of the phage’s physical size. This was important, as previous work had shown alternatively coding phages have genome sizes, and presumably corresponding physical sizes, that range from very small to very large(*4*).

The LC-MS/MS data was searched against phage proteome databases generated by using either code 11 or code 15. Identified peptides were evaluated codon by codon to determine whether translation using standard or alternative genetic code was appropriate. To complement the database search strategy, *de novo* peptide sequencing, which derives peptide sequence information directly from the MS/MS spectra, was incorporated into the traditional database search workflow to provide a database-independent confirmation of phage translation that is agnostic to the translation code used for gene predictions.

Database searching yielded 167 phage-specific peptides in total, with peptide-level false discovery rates at <1%. These peptides mapped to 13 and 14 phage proteins in the infant and adult samples, respectively. In addition, numerous peptides from bacteria and humans were identified (**Data S1**). Many of the phage peptides identified by database searching were further supported by *de novo* sequencing tags. Roughly half of the identified phage peptides in each sample mapped only to proteins predicted using genetic code 15. **Fig. 1** shows the genome maps of the target phages in each sample, with the locations of predicted and detected proteins using either code 11 or code 15 translation. Some of the proteins identified with code 15 predictions were annotated as structural proteins, including capsid, portal, and tail-associated proteins (**Data S1**), while the remaining proteins were unannotated. The detection of mostly structural proteins was expected based on the enrichment for viral-like particles employed for sample preparation.

**Fig. 1.**
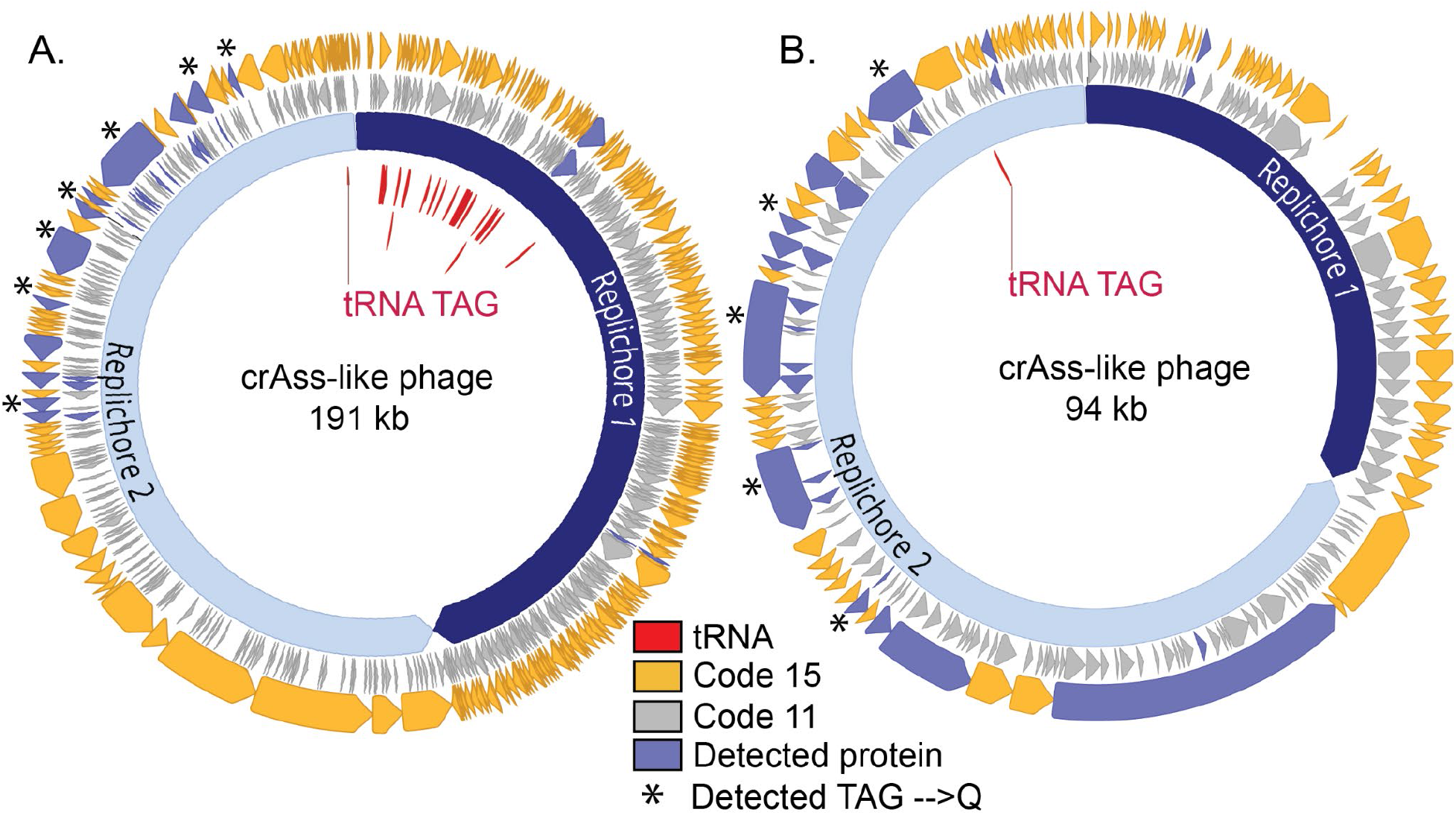
Proteomic detection of alternatively coded proteins from two phage genomes. L2_026_000M1_scaffold_35 (**A**) and L3_063_250G2_scaffold_974_curated (**B**) are alternatively coded crAss-like phages. Prediction of genes in code 11 (inner grey ring) leads to gene fragmentation and low coding density, while gene prediction in code 15 (outer yellow genes) restores open reading frames. Genes with detected peptide evidence are colored purple. Some detected peptides contain glutamines encoded by reassigned TAG codons, and these genes with these validated recoding events are marked with stars. Suppressor tRNAs (red labels) are predicted to suppress translation termination at recoded TAG stop codons. Individual replichores were identified based on GC skew patterns indicative of bidirectional replication.

**Fig. 2** shows the protein sequence coverage map from the alternatively coded phage tail fiber protein (L3_063_250G2_scaffold_974_curated_39.code15) identified in the infant fecal sample. The region of the phage genome corresponding to this single protein contained six truncated proteins when predicted using the standard code. However, when using code 15, the full-length alternatively coded protein contained 23 peptides identified through database matching, of which, 11 were exclusively identified using code 15. Four peptides, highlighted in red boxes, directly confirm that the TAG stop codon is reassigned to glutamine. The identification of several *de novo* sequencing tags provides additional evidence of the existence and expression of recoded stop codons in this alternatively coded protein.

**Fig. 2:**
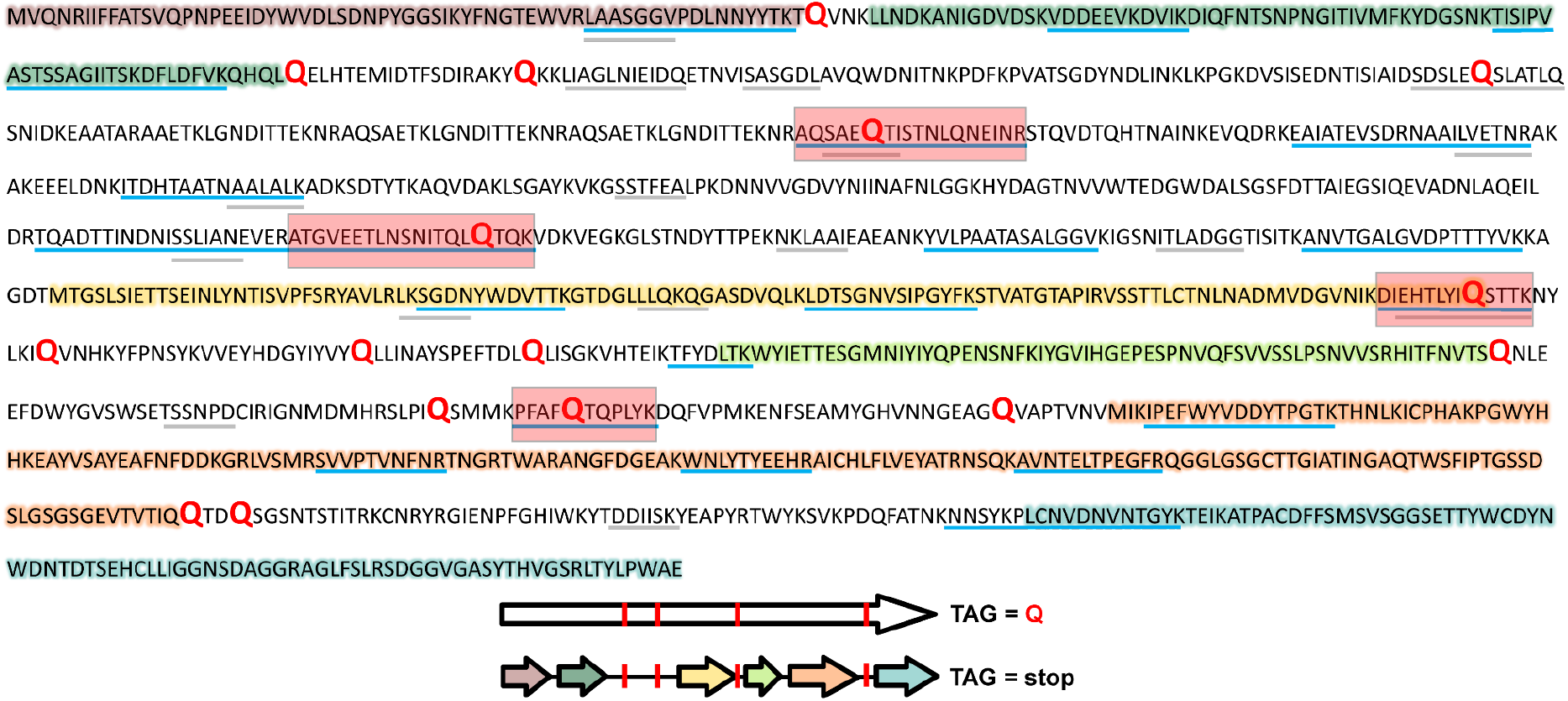
Protein sequence coverage map of alternative code phage tail-related protein. Highlighted sections of the code 15 predicted protein sequence (top) show the corresponding proteins that would have been predicted using standard code 11 (predicted open reading frames), also depicted in the graphical representation (bottom). Blue lines illustrate regions covered by tryptic peptides identified through LC-MS/MS database matching, whereas gray lines represent regions of the predicted protein sequence with matching *de novo* sequence tag coverage. Red text in the sequence indicates the location of glutamine residues from reassigned stop codons. Red boxes on the sequence coverage map and red bars on graphical representation indicate the recoded glutamine residues with peptides identified through database searching.

Numerous identified peptides in both the infant and adult fecal samples further substantiate phage reassignment of the TAG stop codon to glutamine. **Fig. 3** shows two examples of high-quality MS/MS spectra for alternatively coded phage peptides. In both instances, the glutamine residue from the recoded stop codon was positioned in the middle of a tryptic peptide. In the figure, only the direct y-type fragment ion series was chosen for annotation due to their preferential generation in higher-energy C-trap dissociation (HCD) fragmentation during MS/MS measurement(*12*).

**Fig. 3:**
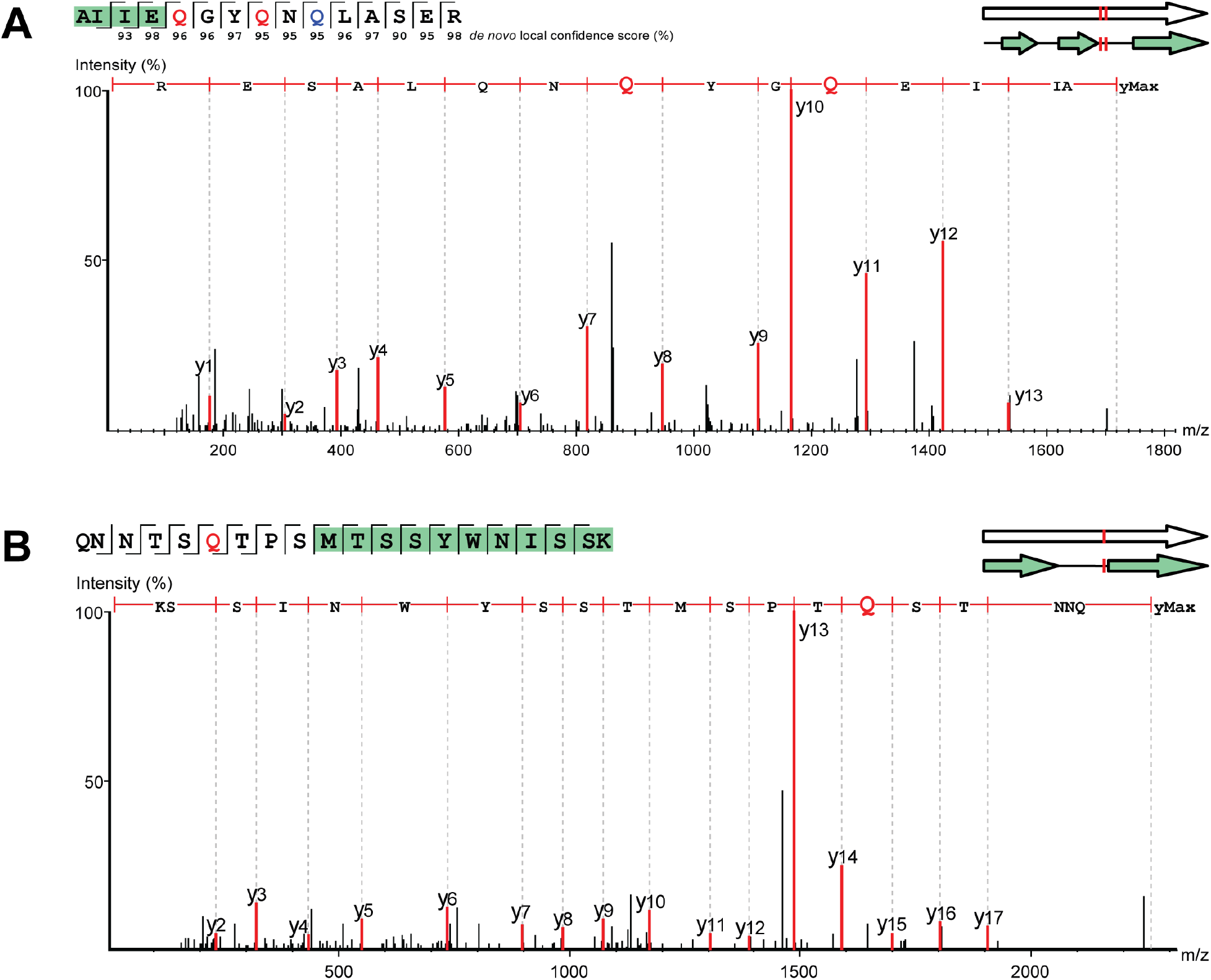
Example MS/MS spectra of alternative coding tryptic peptides. In all panels, red “Q” represents a stop codon that has been reassigned to glutamine using code 15 translation. Blue “Q” represents canonical glutamine residues. Note—the annotated y-ion series are read from the c-terminus to the n-terminus. (top left) Residues highlighted in green on the amino acid sequence fragmentation ladder depicting b-and y-ion fragmentation series represent portions of the peptide also predicted through code 11 (standard code). (**A**) Read-through of a standard genetic code 11 stop codon (L2_026_000M1_scaffold_35_232). (**B**) Read-through of a standard genetic code 11 start codon (L3_063_250G2_scaffold_974_curated_32).

The peptide in **Fig. 3A** contains three glutamine residues; one canonical glutamine and two glutamines from recoded stop codons. One of the recoded glutamines was predicted as a stop codon at the end of a protein predicted through standard code translation. With a nearly complete fragmentation ion series, the detected tryptic peptide shows several amino acids flanking this recoded stop codon, covering an amino acid sequence that would not exist in a standard code open reading frame. In addition, a *de novo* sequencing tag matching nearly the entire length of the database match had high local confidence scores for every amino acid residue, including the recoded glutamines, providing additional support that this peptide, and others like it, do in fact exist (**Fig. S2**). **Fig. 3B** shows a peptide containing a methionine from a predicted start codon using standard code residing in the middle of the peptide sequence in addition to a glutamine from a recoded stop codon. As several amino acids depicted here map to codons upstream of the standard code methionine start codon, this tryptic peptide would not exist if the phage was using standard code translation. Overall, these examples provide experimental validation that standard genetic code 11 is not being utilized by the phage in the translation of this region of the genome.

It has been suggested that alternatively coded structural proteins may be a strategy employed by phages to prevent premature expression of structural and lytic phage genes during the replication process^3^. In this study, the combination of metagenomics and metaproteomics confirmed that when it occurs within genes, the TAG stop codon is translated as glutamine. Direct metaproteomic confirmation of alternative codes has rarely been performed, but it is easy to imagine extending this approach to validate other types of alternative coding, such as the use of alternative start codons or incorporation of non-standard amino acids such as selenocysteine and pyrrolysine. A more complete understanding of how phages modulate the genetic code, likely to ensure appropriate translation of their proteins, may have applications in synthetic biology (e.g., where non-standard reading of codons can be used to create non-biological polymers(*13*)) and in phage engineering.

## Acknowledgments

We acknowledge Dr. Paul Abraham for technical review of this manuscript.

## Funding

National Institutes of Health grant R01GM103600 (J.F.B., R.L.H., M.J.M.)

UC-Berkeley Miller Basic Research Fellowship (A.L.B.)

Oak Ridge Innovation Institute’s EERE Workforce Development project fellowship (S.L.P.)

## Author contributions

Conceptualization: ALB, JFB

Methodology: SLP, RJG, RLH

Investigation: SLP, ALB, RJG, JFB, RLH

Visualization: SLP, ALB, JFB

Funding acquisition: JFB, RLH

Project administration: MJM, JFB, RLH

Supervision: JFB, RLH

Writing – original draft: SLP, ALB, JFB, RLH

Writing – review & editing: SLP, ALB, RJG, MJM, JFB, RLH

## Competing interests

Authors declare that they have no competing interests.

## Data and materials availability

All raw mass spectra and databases for the metaproteome measurement from this study have been deposited into the ProteomeXchange repository with accession numbers: ProteomeXchange-PXD030388; MassIVE-MSV000088561. Genomes are publicly available as an analysis project via ggKbase (https://ggkbase.berkeley.edu/Alternatively_coded_phage_proteomics/organisms).

## Supplementary Materials

Materials and Methods

Figs. S1 to S2

References (*14–23*)

Data S1-S6

## Supplementary Materials

### Materials and Methods

#### Methods

##### Sample selection

In a previous study, human adult and infant stool samples were collected and sequenced with short-read shotgun sequencing (*14*). The adult and infant samples prioritized for proteomics here were chosen because they both had alternatively coded crAss-like phages present at high abundance. Phage L2_026_000M1_scaffold_35 is the most abundant genome in the adult sample at 659x sequencing coverage, and the next highest coverage genome is a *Bacteroides vulgatus* genome at 307x coverage. Phage L3_063_250G2_scaffold_974 is the most abundant genome in the infant sample at 4752x sequencing coverage, and the next highest coverage genome is a *Bacteroides vulgatus* genome at 242x coverage.

##### Genome predictions and phage genome curation

Coding sequences were predicted by Prodigal(*15*) using standard genetic code 11 and alternative genetic code 15. HMMER(*16*) (hmmsearch) was used to annotate protein sequences with the PFAM, pVOG, VOG, and TIGRFAM HMM libraries. In some cases, BLAST searches against the NCBI database, and remote homology searches using the HHPred(*17*) and Phyre2(*18*) web portals were used to augment initial annotations. tRNAs were predicted using tRNAscan-s.e. V.2.0 in general mode(*19*). Replichores were identified by calculating GC skew (G-C/G+C) and cumulative GC skew using the iRep package (gc_skew.py)(*20*). Genome curation was performed using previously described methods(*21*). Curation and genome figure generation were completed using Geneious Prime^®^ 2021.0.3 (https://www.geneious.com/).

##### Sample preparation for LC-MS/MS

100 mg stool sample was resuspended in 1200uL 100mM Tris-HCl, pH 8.0 and homogenized with 0.9-2.0mm stainless steel beads (NextAdvance, part #SSB14B). Homogenized samples were incubated for 60 minutes before centrifugation at 3,000xg for 30 minutes. After centrifugation, the pre-cleared supernatant was filtered on a 300kDa MWCO PES filter (Pall, Omega Membrane 300K, part # OD300C34). The filtered eluate and the residual proteinaceous biomass remaining on top of the filter (resuspended in 100mM Tris-HCl, pH 8.0) were collected for downstream processing. Samples were adjusted to 4% (wt:wt) sodium dodecyl sulfate (SDS)/5mM dithiothreitol and incubated at 95°C for 10 min. Samples were alkylated with 15mM iodoacetamide (IAA) for twenty minutes at room temperature in the dark. The crude protein sample volume was processed by the protein aggregation capture (PAC) method(*22*). Briefly, 300ug of magnetic beads (1 micron, SpeedBead Magnetic Carboxylate; GE Healthcare UK) was added to each sample. Samples were then adjusted to 70% acetonitrile to induce protein aggregation. Aggregated proteins were washed on a magnetic rack with 1mL of 100% acetonitrile, followed by 1mL of 70% ethanol. The washed proteins were then resuspended in 4% SDS/ 100mM Tris-HCl, pH 8.0, and boiled off the magnetic beads at 95°C for 10 minutes. Protein amounts were quantified by corrected absorbance (Scopes) at 205 nm (NanoDrop OneC; Thermo Fisher). The cleaned proteins were re-aggregated back on the magnetic beads and washed with 1mL of 100% acetonitrile and 1mL of 70% ethanol to remove detergent from samples. The aggregated protein pellet was resuspended in 100mM Tris-HCl, pH 8.0, and digested with 1:75 (wt:wt) proteomics-grade trypsin (Pierce) overnight at 37°C and again for four hours the following day. Tryptic peptides were filtered on a 10kDa MWCO filter plate (AcroPrep Advance, Omega 10K MWCO) at 12,000xg and adjusted to 0.5% formic acid before quantification by NanoDrop OneC.

##### LC-MS-MS

Digested peptides were analyzed by automated 1D LC-MS/MS analysis using a Vanquish ultra-HPLC (UHPLC) system plumbed directly in line with a Q Exactive Plus mass spectrometer (Thermo Scientific). A trapping column (100 μm inner diameter; packed with 5 μm Kinetex C18 reverse-phase resin (Phenomenex) packed to 10 cm) was coupled to an in-house-pulled nanospray emitter (75 μm inner diameter; 1.7 μm Kinetex C18 reverse-phase resin (Phenomenex) packed to 15 cm). For each sample, 10uL of peptides were loaded, desalted, and separated by uHPLC under the following conditions: sample injection followed by 100% solvent A (95% H2O, 5% acetonitrile, 0.1% formic acid) from 0-30 minutes to load and desalt, a linear gradient from 0% to 30% solvent B (70% acetonitrile, 30% water, 0.1% formic acid) from 30-220 minutes for separation, and 100% solvent A from 220-240 minutes for column re-equilibration. Eluting peptides were measured and sequenced with a Q Exactive Plus MS under the following settings: data-dependent acquisition; mass range 300 to 1,500 m/z; MS and MS/MS resolution 70K and 15K, respectively; MS/MS loop count 20; isolation window 1.8 m/z; charge exclusion unassigned, 1, 6 to 8.

##### Proteomics data analysis

MS/MS spectra were interrogated by *de novo*–assisted database searching(*23*) in PEAKS Studio 10.6 (Bioinformatics Solutions) against a custom-built proteome database derived from the combination of the sequenced metagenome-derived predicted proteomes for all contig except the target phage contig, the phage proteome predicted in standard code (code 11), and the phage proteome predicted in the alternative code (code 15), the human reference proteome from UniProt (UP000005640), common LC-MS/MS protein contaminants, and reversed-decoy sequences of all proteins in the database. The parent and fragment ion mass error tolerances were set to ± 10 ppm and ±0.02 Da, respectively. Peptide spectrum matches (PSMs) were required to be tryptic with semi-specific digestion needed and a maximum of three missed cleavages. Accepted modifications included a fixed modification of carbamidomethylation (+57.02) of cysteine residues and a variable modification of methionine oxidation (+15.99), with a maximum of three variable modifications. A false discovery rate of 1% was applied to accept the peptide and protein sequences, and a minimum of one unique peptide was required to identify a protein. *De novo* only parameters were left at default settings with average local confidence (ALC) scores of >50% and *de novo* sequence tags displayed if at least six amino acids were shared with the database sequence. The resulting database-identified peptides and corresponding *de novo* sequence tags were manually validated to generate the final list of phage peptide sequences present in the sample. A database hit passing the 1% FDR threshold was required for a peptide sequence to be considered as detected. *De novo* sequence tags were regarded as complementary evidence for the code 15 database-identified peptides to confirm the expression of this code only if the residue local confidence scores of the amino acids in the *de novo* sequence tag matching the database sequence were greater than 90% confidence for each residue. In the cases where a TAG stop codon was reassigned to glutamine, the glutamine and several flanking amino acids in the *de novo* sequence tag had to pass the >90% residue local confidence score threshold.

**Fig. S1.**
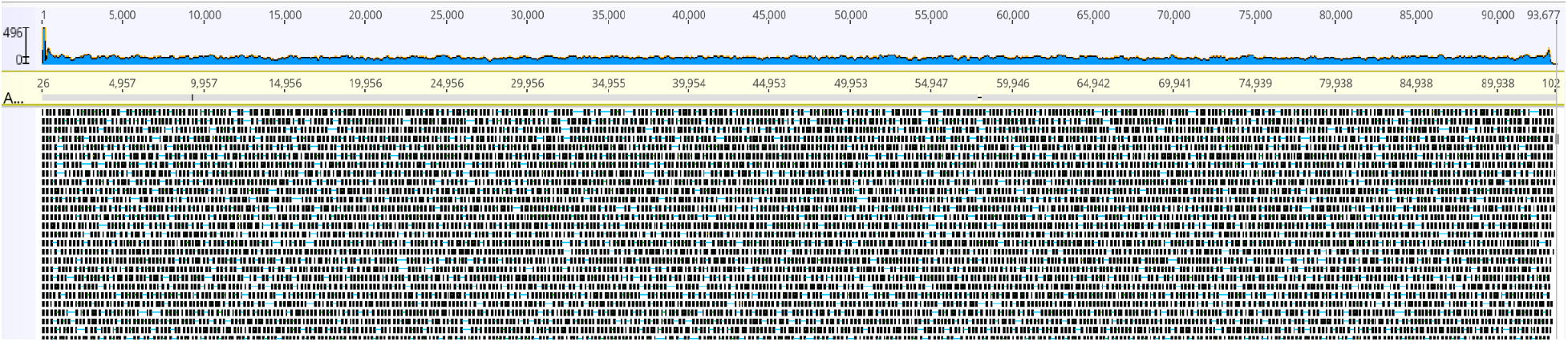
Genome curation and variation. Overview of the final curated genome showing complete and relatively even coverage by paired 150 bp Illumina reads (mean insert size: 391 bp) when reads are mapped, allowing ≤ 2% single nucleotide polymorphisms.

**Fig. S2.**
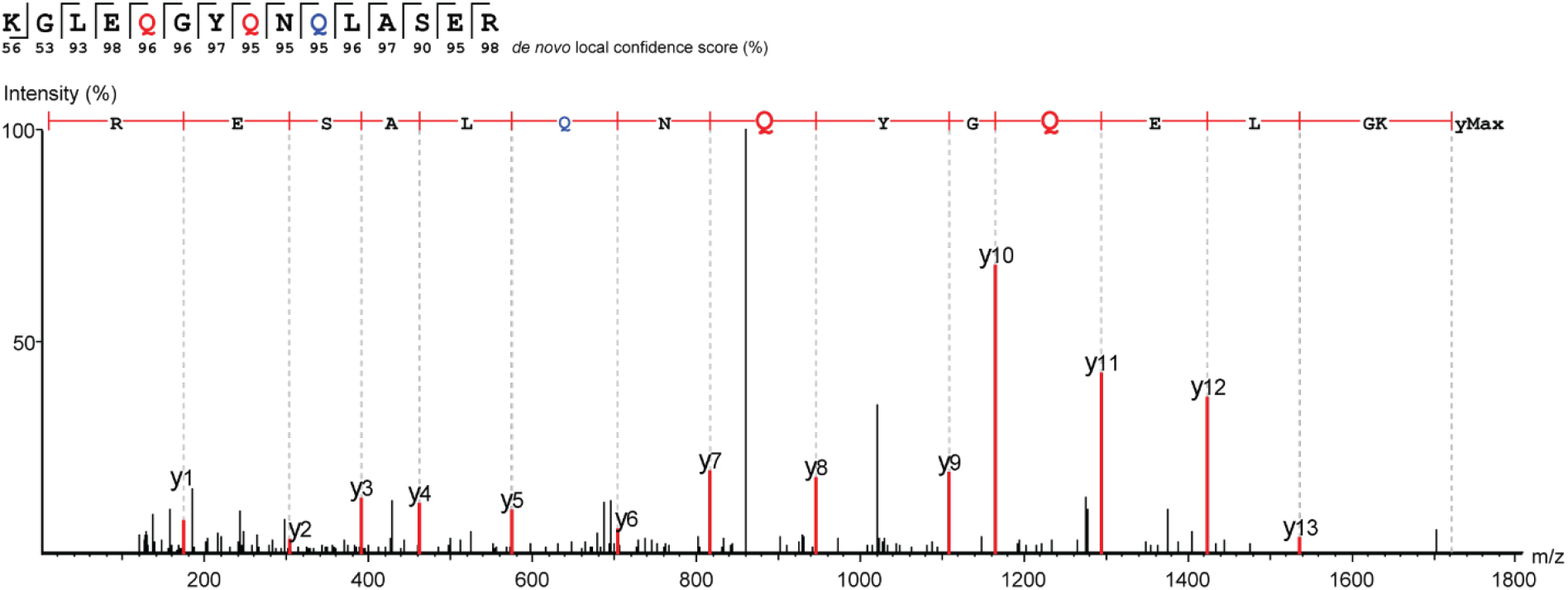
MS/MS spectrum of *de novo* sequence tag of an alternative coded tryptic peptide. The de novo sequence tag matches nearly the entire amino acid sequence of the database search identified peptide in Fig. 3A. The canonical glutamine and both glutamine residues from recoded stop codons, as well as flanking residues, have very high *de novo* local confidence scores.

**Data S1. (separate file)**

**L2_026_000M1 detected peptides.** This table contains all peptides (bacterial, phage, human) from fecal sample L2_026_000M1 identified by LC-MS/MS with a peptide-level false discovery rate (FDR) threshold of 1%.

**Data S2. (separate file)**

**L2_026_000M1 detected proteins.** This table contains all proteins (bacterial, phage, human) from fecal sample L2_026_000M1 identified by LC-MS/MS with a protein-level false discovery rate (FDR) threshold of 1% and at least one unique peptide per protein.

**Data S3. (separate file)**

**L2_026_000M1_scaffold_35 detected proteins and peptides**. This table contains all detected phage proteins from scaffold L2_026_000M1_scaffold_35. For each protein detected using code 15 predictions (dark green), any corresponding protein predicted using code 11 (light green) with peptide evidence is listed below the code 15 protein. Nested rows below each protein correspond to the peptide analytes detected for the protein. The column “#Peptide analytes” refers to both unmodified and modified versions of the peptide sequence. Peptide rows are colored to show whether the peptide was found using both code 11 and code 15 predictions (light yellow) or only through code 15 prediction (dark yellow). Red text indicates peptides with at least one sequenced glutamine from a recoded stop codon.

**Data S4. (separate file)**

**L3_063_250G2 detected peptides**. This table contains all peptides (bacterial, phage, human) from fecal sample L3_063_250G2 identified by LC-MS/MS with a peptide-level false discovery rate (FDR) threshold of 1%.

**Data S5. (separate file)**

**L3_063_250G2 detected proteins.** This table contains all proteins (bacterial, phage, human) from fecal sample L3_063_250G2 identified by LC-MS/MS with a protein-level false discovery rate (FDR) threshold of 1% and at least one unique peptide per protein.

**Data S6. (separate file)**

**L3_063_250G2_scaffold_974 detected proteins and peptides**. This table contains all detected phage proteins from scaffold L3_063_250G2_scaffold_974. For each protein detected using code 15 predictions (dark green), any corresponding protein predicted using code 11 (light green) with peptide evidence is listed below the code 15 protein. Nested rows below each protein correspond to the peptide analytes detected for the code 15 protein. The column “#Peptide analytes” refers to both unmodified and modified versions of the peptide sequence. Peptide rows are colored to show whether the peptide was found using both code 11 and code 15 predictions (light yellow) or only through code 15 prediction (dark yellow). Red text indicates peptides with at least one sequenced glutamine from a recoded stop codon.

## Notes

https://ggkbase.berkeley.edu/Alternatively_coded_phage_proteomics/organisms

### Competing Interest Statement

The authors have declared no competing interest.

